# Mediterranean oaks harbor more specific soil microbes at the dry end of a precipitation gradient

**DOI:** 10.1101/2020.05.14.095943

**Authors:** Gemma Rutten, Lorena Gómez-Aparicio, Beat Frey

## Abstract

**Background:** Recent evidence suggests that soil microbial communities can regulate plant community dynamics. In addition, the drought tolerance of plants can be enhanced by soil microbes. So far, few studies have assessed the variation in the microbiome of specific plant species along environmental gradients. Yet understanding these dynamics is essential to improve predictions of plant-soil feedbacks and the consequences of ongoing climate changes. Here we characterized the soil microbiome of two co-occurring Mediterranean oaks along a precipitation gradient, using amplicon sequencing of phylogenetic marker genes for prokaryotes and fungi. Additionally, we identified tree-specific and locally-specific microbes potentially responsible for tree community dynamics.

**Results:** We show that two co-occurring, evergreen Mediterranean oak species harbor distinct microbiomes along a precipitation gradient. The soil microbial diversity increased along the precipitation gradient, for prokaryotic α and β diversity and for fungal β diversity. *Quercus ilex* harbored richer fungal communities than *Quercus suber*, and host-specific taxa more often belonged to fungi than to prokaryotes. Notably, the microbial communities at the dry end of the precipitation gradient harbored more locally-specific prokaryotic and fungal taxa than the microbial communities with a higher diversity, at the wet end of the gradient, suggesting higher specialization in drier areas.

**Conclusions:** Even congeneric tree species, belonging to the same functional group, can harbor distinct and specific soil microbiomes. These microbiomes become more similar and consist of more specialized taxa under drier compared with wetter conditions. With this, our study offers a step towards a better understanding of the context-dependency of plant-soil feedbacks by going beyond α and β diversities and focusing on specialized taxa potentially driving community changes along environmental gradients. We hope that our study will stimulate future research assessing the importance of context-dependency of interactions between plants and soil communities in a changing world.

## Introduction

Interactions between plants and microbes affect ecosystem functioning [1–3], and plant-soil feedbacks (PSFs) have been shown to be relevant drivers of forest community dynamics [4,5]. Moreover, specific root-associated microbes have been found to enhance the drought tolerance and overall health of plants [6–8]. Few studies however have explored the variation in plant-microbiome interactions along environmental gradients, yet understanding these dynamics is essential for improving predictions of the consequences of ongoing climate changes.

At the global and regional scales, precipitation is among the most important drivers of soil microbial diversity [9–12], together with other variables like temperature [13,14] and soil pH [15–18]. However, the diverse life history strategies of different taxa can result in numerous responses along environmental gradients [18]. Reduced water availability and increased drought intensity lead to reduced microbial growth and shifts in microbial community composition [19,20]. In particular, decomposers, which contribute to the carbon cycle, have been found to have a lower activity in drier sites than in wetter sites [21]. Food webs with a fungal basis are generally more resistant to drier conditions than bacteria-based food webs [22] but small changes in water availability can result in shifts in fungal species dominance, whereas bacterial communities remain constant [23]. Repeated wet-dry cycles can result in adapted soil communities that show fewer shifts in community composition with fluctuations in water availability [24,25].

Soil microbial communities are not only shaped by abiotic factors such as precipitation and water availability, also biotic drivers such as plants play an important role. Plants can affect microbial communities both directly and indirectly. For example, mycorrhizal fungi and fungal pathogens can accumulate directly in the rhizosphere of the host plant. Moreover, species-specific litter quality and root exudates might indirectly affect the microbial community through changes in nutrient availability and soil properties [26,27]. Prokaryotes are more likely related to differences in soil properties [28], potentially partly caused by plants, even though they can also be plant species-specific [7,29]. Most existing studies have focused on associations between plants and fungi, as fungi are more host-specific than prokaryotes [30].

Under dry conditions, communities are often dominated by highly specialized species [31] and the interactions among species become more facilitative as predicted by the stress gradient hypothesis [32,33]. For instance, under dry conditions, species-specific microbes can mitigate water stress for their hosts [6,7,29]. Particularly mycorrhiza can help increase the resource uptake of plants and can defend a plant against pathogens [34]. Therefore, both mutualists and pathogens can play an important role in facilitative interactions [35]. Conspecific seedlings experience more positive effects from the soil microbial community when growing in drier than in wetter conditions, suggesting more beneficial effects of the soil community under drier conditions [3,36,37]. However, if specialized mutualists or pathogens are responsible for these shifts along environmental gradients remains largely untested [6]. To disentangle how plant-microbiome interactions vary along climatic gradients, we need to combine the microbial perspective, often focusing on variations in nutrient cycling and decomposition, with the plant perspective, focusing mainly on the mutualistic and antagonistic soil community components.

In this study, we simultaneously tested the effects of precipitation and tree species on soil microbial communities in a Mediterranean climate. Specifically, we used high throughput amplicon sequencing to characterize the prokaryotic and fungal communities under the two dominant evergreen oak species in southern Spain, *Quercus ilex* and *Quercus suber*, along six sites ranging in precipitation (637, 672, 751, 890, 912, 948 mm year^−1^). We expected 1) a reduced microbial community richness and diversity in areas with low precipitation, particularly for prokaryotes. Additionally, we expected 2) stronger effects of tree species on fungal than on prokaryotic community diversity. Moreover, we tested 3) the effects of local variations in soil properties to account for indirect effects of trees on soil microbial communities. Concerning the specialized soil community components, we expected 4) more fungal than prokaryotic taxa to be tree-specific. Finally, following the stress gradient hypothesis, we expected 5) more locally-specific microbes in drier areas and more generalist in more humid areas. Testing these hypotheses will help to better understand the variation in plant-microbiome interactions along environmental gradients and ultimately improve predictions of the consequences of ongoing climate changes.

## Methods

### Experimental design and site selection

We used the third Spanish National Forest Inventory (NFI3; Ministry of Environment, Madrid 2006) to select study sites where the two most abundant evergreen Mediterranean oak species in the country (*Quercus suber* and *Quercus ilex*) co-occur. In November 2015, we selected six sites covering a precipitation gradient (637, 672, 751, 890, 912, 948 mm year^−1^) in south-central Spain [36]. The sites were located between 100 and 594 m a.s.l and had mean annual temperatures between 15°C and 18°C. Each of the six sites contained four *Q. ilex* and four *Q. suber* adult trees (48 individual trees in total), which had no woody understory or overlapping crowns. In January 2016, we collected soil samples (0-15 cm depth) from under the crown of each tree using a soil corer (5 cm diameter). We pooled 9 to 15 cores to generate one composite soil sample per tree. The 48 pooled soil samples were kept in a cooling container until further processing in the lab.

### Physical and chemical soil properties

In the lab, the soil samples were homogenized, sieved through a 2 mm sieve to remove roots and mesofauna, and split into two parts. One part was air dried for three days 25°C and subsequently analyzed for soil physical and chemical properties according to standardized protocols [58]. As we could not separate indirect tree species effects, through leaf litter input, nutrient depletion and root exudation, from inherent site differences, we assessed physical and chemical soil properties as a whole. In short, we assessed soil texture, pH, total soil organic matter (SOM) content, and concentrations of nitrate (NO_3_^-^), ammonium (NH_4_^+^), available phosphorus (P), potassium (K), calcium (Ca) and magnesium (Mg). For additional information see Supplement 1.

### DNA extraction, PCR amplification and sequencing

We used the second part of the soil sample (0.25 g) to extract DNA using the PowerSoil isolation kit (Mo-Bio Laboratories Inc., Carlsbad, CA, USA) according to the manufacturer’s instructions. Extracted DNA was quantified using Picogreen (Molecular Probes, Eugene, OR, USA). We amplified the partial prokaryotic small-subunit ribosomal RNA genes (region V3–V4 of 16S rRNA gene) and the fungal ribosomal internal transcribed spacers (region ITS2), as described previously [59]. Each sample containing 40 ng of DNA was amplified in triplicate and pooled before purification with Agencourt AMPure XP beads (Beckman Colter, Berea, CA, USA). The amplicons were sent to the Genome Quebec Innovation Centre (Montreal, Canada) for barcoding using Fluidigm Access Array technology and paired-end sequencing on the Illumina MiSeq v3 platform (Illumina Inc., San Diego, CA, USA).

### Bioinformatics

For the quality control of prokaryotic and fungal reads, we used a customized pipeline [59]. Briefly, we matched paired-end reads [60] correcting for substitution errors [61], and trimmed the PCR primers, with one mismatch allowed [62]. We discarded prokaryotic sequences (16S rRNA gene, region V3V4) shorter than 300 bp, fungal sequences (region ITS2) shorter than 200 bp, and sequences without matching primers. The fastq_filter function in USEARCH was used for quality filtering of the trimmed reads with a maximum expected error threshold of one. All singletons were removed to reduce artificial OTU inflation while clustering [63]. We used the cluster_otu function in USEARCH, which includes an ‘on-the-fly’ chimera removal algorithm to cluster the sequences in OTUs applying a 97% similarity threshold. Metaxa2 [64] and ITSx [65] were used to test the centroid OTU sequences for the presence of ribosomal signatures, and unsupported centroid sequences were discarded. Remaining reads that passed the quality filtering, were mapped with USEARCH usearch_global algorithm (parameters: maxrejects 0, maxaccepts 0 and top_hit_only) against the final OTU list. All raw sequences were deposited in the NCBI Sequence Read Archive under the BioProject accession number PRJNA550539. We used MOTHUR with its implemented naïve Bayesian classifier (minimum bootstrap support 60%) to query the centroid sequences to the reference databases. SILVA Release 128 served as the reference database for the prokaryotic sequences [66]. Eukaryotic sequences were queried against the two databases in NCBI GenBank [67] and UNITE 8.0 [68]. We removed eukaryote sequences (0.6%) and unknown sequences (1.4%) from the prokaryotic OTUs, as well as OTUs not belonging to the fungal kingdom, including plants (2.2%), protists (0.4%) and unclassified eukaryotes (0.9%). To increase the probability of assessing ecologically relevant taxa [69], we pruned the prokaryotic and fungal datasets of rare OTUs (total abundance <10 reads; occurring >3 samples).

### Statistical analysis

To test for differences in alpha diversity between the oak species along the precipitation gradient we first made rarefaction curves. These curves visualize sampling effort and help determine the sample with the smallest number of reads (Figure S1). To correct for increasing species numbers with an increasing number of reads, we rarefied the data by the smallest total number of sequences per sample, resulting in 15,548 prokaryotic and 12,740 fungal reads per sample. This step led to the removal of 10 fungal OTUs from the data. Of the many indexes available to quantify alpha diversity, we chose to calculate the observed richness (S_obs_), as it is widely used and easily comparable between studies. We calculated other alpha diversity measures for each sample, for both raw and rarified datasets [69], and found small differences depending on the measure used (data not shown). To test for the effects of precipitation and tree species on the diversity indexes, we used generalized linear mixed effects models (GLMM) using *lme4* [70], and included site as a random factor to control for multiple samples coming from the same site. In addition, we calculated the relative abundance of OTU counts for each phylum (with a total abundance >1%) and for each treatment combination. Finally, we assessed the effects of precipitation and tree species on the relative abundance of the microbial sequences, using PERMANOVAs based on Euclidian distances with 9999 permutations, using *vegan* [71]. To account for site effects, we included this factor in the random effects (strata).

To quantify differences in community structure (beta diversity) across the precipitation gradient and between tree species, we used PERMANOVAs based on Bray-Curtis distances with 9999 permutations, with site included in the strata. The Bray-Curtis dissimilarity matrices were generated based on either relative or normalized OTU sequence counts of the unrarefied data. To normalize the data, we performed a transformation by the weighted trimmed mean of M-values [72,73]. Applying alternative transformations [69,74,75] yielded similar results and are therefore only presented in the supplementary (Table S1). Differences in the community structure among sites and between tree species were visualized with unconstrained PCoA-ordinations and with site- and tree-species constrained CAP-ordinations [71]. To determine which edaphic variables (pH, SOM, NO_3_^-^, NH_4+_, P, K, Ca, Mg and proportion of sand (as a measure of soil texture) significantly explained variation in the prokaryotic and fungal communities, we used non-metric multidimensional scaling (NMDS) in *vegan* [71].

To identify which OTUs led to changes in multivariate patterns between tree species and along the precipitation gradient, we studied the relationship between OTUs and groups of sites using indicator species analysis [76,77]. First, we investigated which taxa were significantly associated to a tree species, hereafter referred to as species-specific taxa. We combined an indicator species analysis with GLMMs and log-likelihood-ratio tests implemented in the *EdgeR:glmLRT* and *EdgeR:topTag* [72,73]. To explore if more locally-specific taxa exist in drier sites, we tested which taxa in the prokaryotic and the fungal communities were more abundant than expected based on random distributions over the sites [38]. Based on this analysis, we calculated the proportion of locally-specific taxa for each site along the precipitation gradient, for prokaryotes and fungi separately. We then used a linear model, with precipitation and tree species as fixed factors, to test whether the proportion of locally-specific taxa decreased along the precipitation gradient. All statistics were performed in R [78] using the cited packages, functions and their dependencies.

## Results

### Diversity of prokaryotes and fungi under two oak species along a precipitation gradient

The α diversity was higher for prokaryotes than for fungi and differed across our treatments (Table 1, Figure 1). For prokaryotes, an increased precipitation tended to result in in more diverse communities, whereas their alpha diversity did not differ among tree species (Table 1, Figure 1a). For fungi, a higher precipitation did not affect the α diversity, but more diverse communities were found under *Q. ilex* than under *Q. suber* (Table 1, Figure 1b). Specifically, the relative abundance of *Ascomycota, Basidiomycota, Verrucomicrobia* and *Gammaproteobacteria* showed significant tree species effects (Table S1, Figure S1), whereas *Mortierellomycota, Acidobacteria* and *Patescibacteria* showed significant interactions between tree species and the precipitation gradient (Table S1, Figure S1).

**Table 1.**
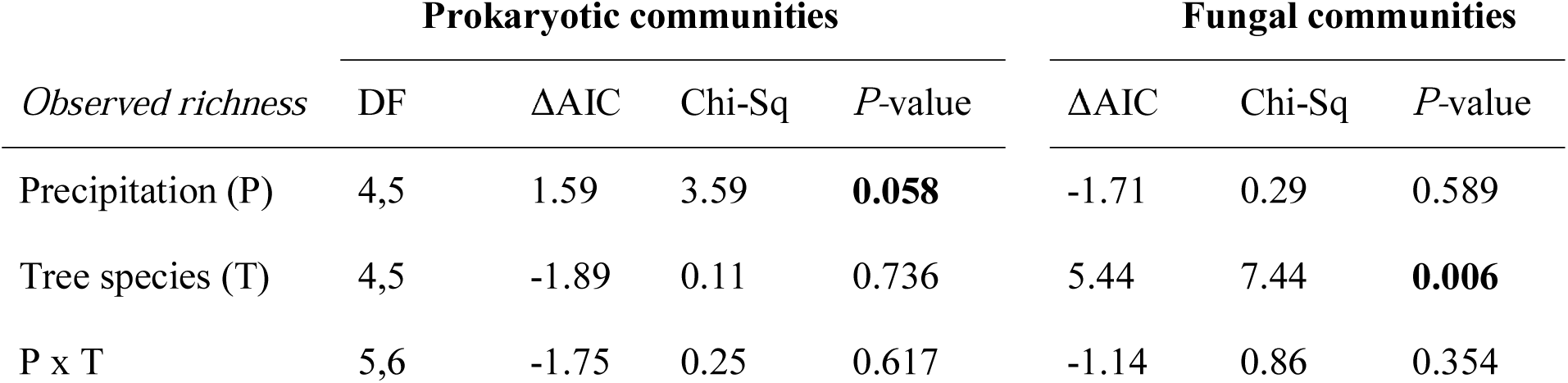
Alpha diversity of the prokaryotic and fungal communities, measured as Observed Richness (S_obs_). Linear models were performed on rarefied community data, with precipitation (P), tree species (T) and their interaction (P x T) as fixed effects and site as a random factor. Significant terms (*P* <0.1) are shown in bold.

| *Observed richness* | **Prokaryotic communities** |  |  |  | **Fungal communities** |  |  |
| --- | --- | --- | --- | --- | --- | --- | --- |
| | DF | $\Delta\text{AIC}$ | Chi-Sq | *P*-value | $\Delta\text{AIC}$ | Chi-Sq | *P*-value |
| Precipitation (P) | 4,5 | 1.59 | 3.59 | **0.058** | -1.71 | 0.29 | 0.589 |
| Tree species (T) | 4,5 | -1.89 | 0.11 | 0.736 | 5.44 | 7.44 | **0.006** |
| P x T | 5,6 | -1.75 | 0.25 | 0.617 | -1.14 | 0.86 | 0.354 |

**Figure 1.**
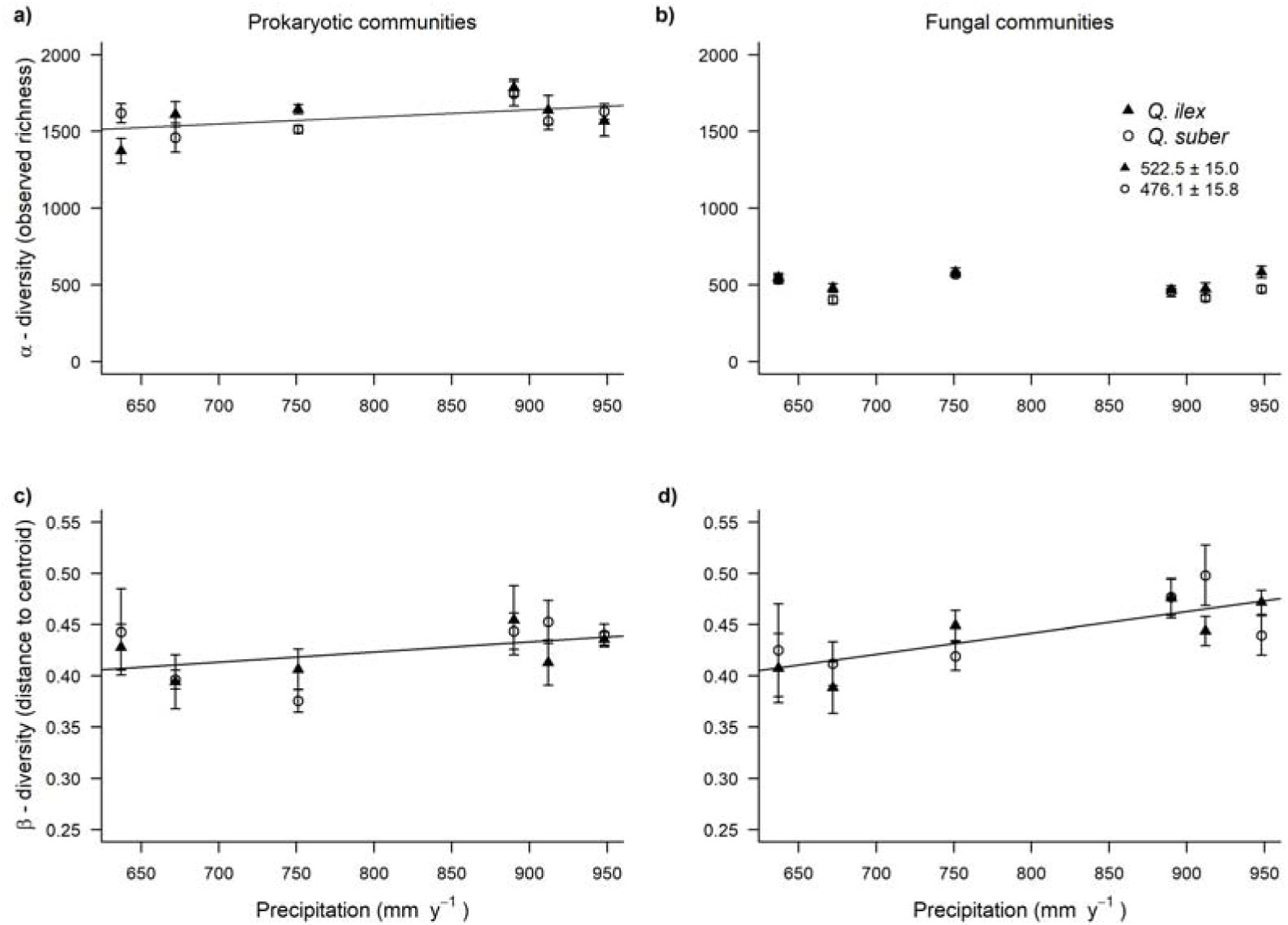
Diversity of microbial communities under *Quercus ilex* trees (triangles) and *Quercus suber* trees (circles) along a precipitation gradient. Alpha diversity (a, b) measured as observed richness of rarefied a) prokaryotic and b) fungal OTUs. Beta diversity (c, d) measured as Euclidean distance in principle coordinate between a sample and the group centroid for c) prokaryotic and d) fungal communities. Dissimilarity indices were calculated using Bray-Curtis distances of relative abundance per sample.

The β diversity of fungal and prokaryotic communities changed along the precipitation gradient (Table 2), with both prokaryotic (Figure 1c) and fungal (Figure 1d) communities becoming more dissimilar with increasing precipitation. Fungal communities were generally more variable whereas prokaryotic communities showed more overlap along the precipitation gradient and between tree species (Figure 1, Figure 2, Figure S2). Additionally, significant interactions between precipitation and tree species for prokaryotic (P x T interaction, *P* = 0.05, Table 2) and fungal communities (P x T interaction *P* = <0.001, Table 2), indicate that the dissimilarity of the soil microbiome under the two tree species is determined by the precipitation regime. Moreover, a significant tree species effect on the β diversity of fungi (*P* = <0.001, Table 2) indicates that the fungal communities under *Q. ilex* were less dissimilar than those under *Q. suber* (Table2; Figure 1).

**Table 2.**
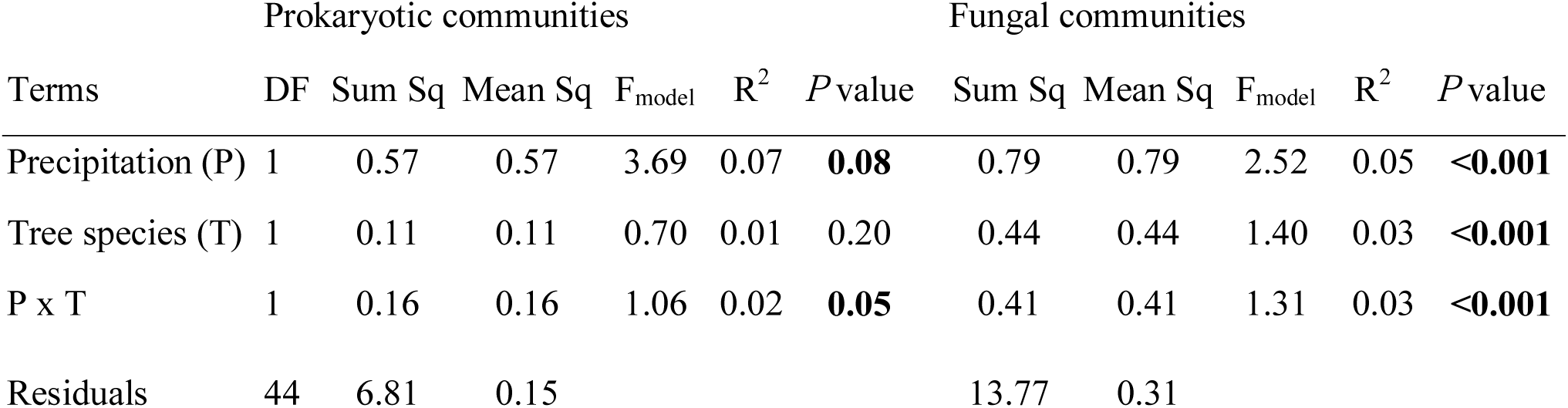
Beta diversity of prokaryotic and fungal communities. PERMANOVAs were performed on *edgeR* normalized data, with precipitation (P) and tree species (T) and their interactions as fixed factors. All models were run with 9999 permutations and site included in strata to correct for site effects. Significant terms (*P* <0.1) are shown in bold.

|  | Prokaryotic communities |  |  |  |  |  | Fungal communities |  |  |  |  |
| --- | --- | --- | --- | --- | --- | --- | --- | --- | --- | --- | --- |
| Terms | DF | Sum Sq | Mean Sq | F<sub>model</sub> | R<sup>2</sup> | P value | Sum Sq | Mean Sq | F<sub>model</sub> | R<sup>2</sup> | P value |
| Precipitation (P) | 1 | 0.57 | 0.57 | 3.69 | 0.07 | **0.08** | 0.79 | 0.79 | 2.52 | 0.05 | **<0.001** |
| Tree species (T) | 1 | 0.11 | 0.11 | 0.70 | 0.01 | 0.20 | 0.44 | 0.44 | 1.40 | 0.03 | **<0.001** |
| P x T | 1 | 0.16 | 0.16 | 1.06 | 0.02 | **0.05** | 0.41 | 0.41 | 1.31 | 0.03 | **<0.001** |
| Residuals | 44 | 6.81 | 0.15 |  |  |  | 13.77 | 0.31 |  |  |  |

**Figure 2.**
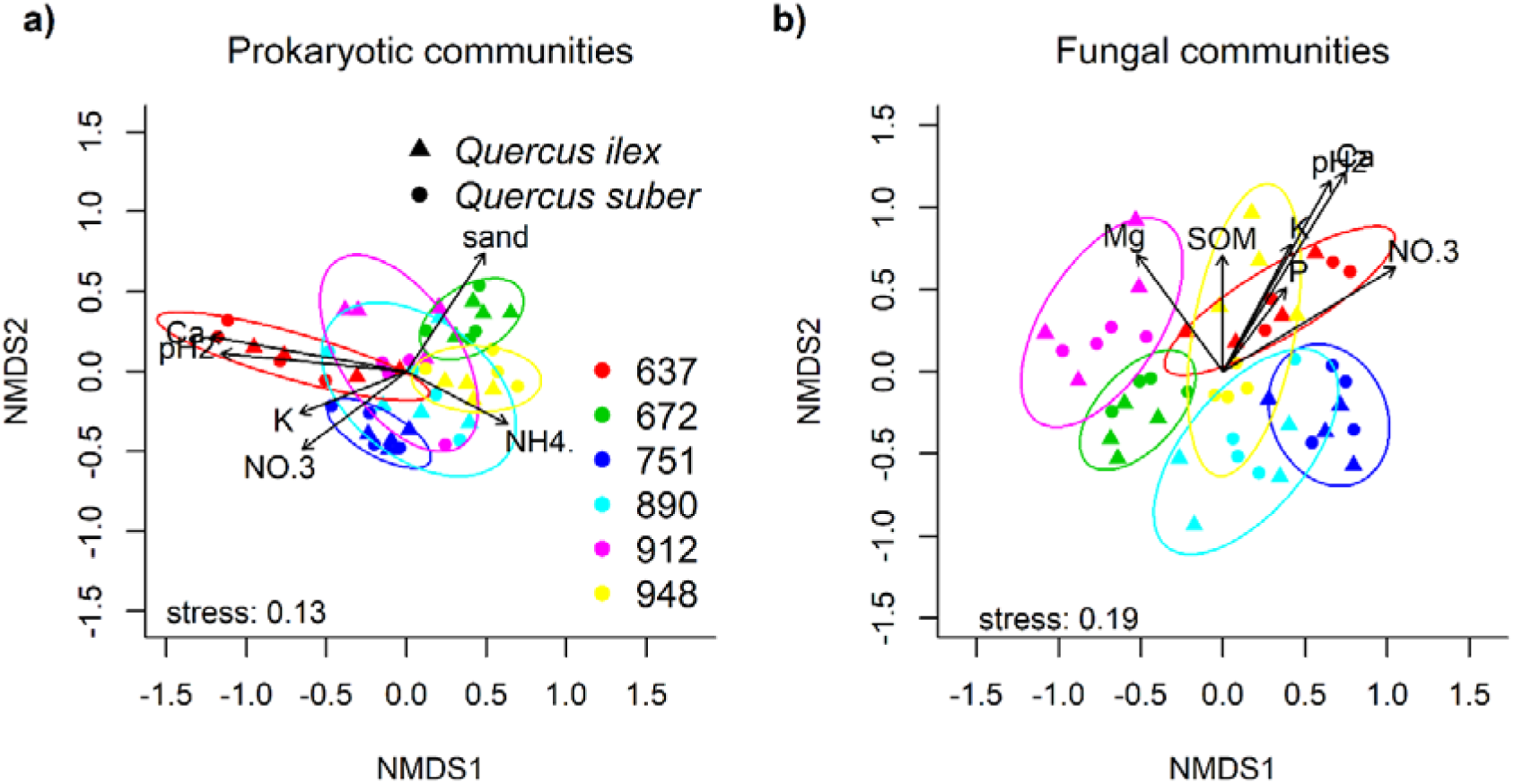
Non-metric multi-dimensional scaling (NMDS) based on the Bray-Curtis distances. Community structure was based on the squared-root relative abundances of (**a**) prokaryotic OTUs and (**b**) fungal OTUs, along the precipitation gradient (indicated by the different colors) and between *Quercus ilex* trees (triangles) and *Quercus suber* trees (circles). Stress values indicate the goodness of fit. Vectors indicate the strength and direction of significant environmental variables (*P*<0.01).

The most influential edaphic conditions on prokaryotic and fungal communities were calcium (explaining 57% of prokaryotic structure and 60% of fungal structure), followed by pH (50% and 53%), and nitrate (24% and 45%), potassium (19% and 24%) and finally phosphorus (10% and 12, Table 3). Additionally, sand and ammonium explained 29% and 19% of the variation in prokaryotic communities, respectively (Table 3, Figure 2), whereas these variables did not affect fungal communities. Conversely, magnesium and SOM explained 23% and 15 % of the variation in fungal communities, respectively, whereas these variables did not impact the prokaryotic communities (Table 3, Figure 2).

**Table 3.** Non-metric Multi-Dimensional Scaling (NMDS) of prokaryotic and fungal communities. Significant correlations between measured soil properties and NMDS are shown in bold and ordered by decreasing explanatory power, measured as R^2^. Soil properties significantly explaining variation in both prokaryotic and fungal communities are shown in italics. Significant terms (*P* <0.1) are shown in bold.

| Vectors | Prokaryotic communities |  |  |  | Fungal communities |  |  |  |
| --- | --- | --- | --- | --- | --- | --- | --- | --- |
| | NMDS1 | NMDS2 | $R^2$ | $P$ value | NMDS1 | NMDS2 | $R^2$ | $P$ value |
| ***Calcium*** | -0.98 | 0.18 | 0.57 | **0.001** | 0.52 | 0.85 | 0.60 | **0.001** |
| ***pH*** | -1.00 | 0.10 | 0.50 | **0.001** | 0.49 | 0.87 | 0.53 | **0.001** |
| ***Nitrate ($NO_3^-$)*** | -0.80 | -0.60 | 0.24 | **0.004** | 0.86 | 0.52 | 0.45 | **0.001** |
| ***Potassium (K)*** | -0.93 | -0.36 | 0.19 | **0.011** | 0.47 | 0.88 | 0.23 | **0.004** |
| **Texture (% sand)** | 0.55 | 0.83 | 0.29 | **0.001** | -0.99 | 0.17 | 0.07 | 0.191 |
| Magnesium (Mg) | -0.98 | -0.18 | 0.03 | 0.445 | -0.59 | 0.81 | 0.23 | **0.006** |
| ***Phosphorus (P)*** | 0.65 | -0.76 | 0.10 | **0.076** | 0.60 | 0.80 | 0.12 | **0.043** |
| SOM | 0.99 | -0.14 | 0.06 | 0.252 | -0.01 | 1.00 | 0.15 | **0.023** |
| **Ammonium ($NH_4^+$)** | 0.89 | -0.46 | 0.19 | **0.008** | 0.72 | 0.70 | 0.02 | 0.665 |

### Specialization of prokaryotes and fungi under two oak species along a precipitation gradient

There was a clear difference in the taxa frequency distribution between the prokaryotic and fungal communities (Figure 3). Only a small proportion of taxa was found in all samples (Figure 3), namely 2.51% (115 OTUs) for prokaryotes and 0.65% (20 OTUs) for fungi respectively. These ubiquitous fungi taxa, included nine saprotrophs, six unassigned guilds, two animal pathogens, two plant pathogens (*Mycosphaerella tassiana* and *Trichoderma asperelluman*) and one endophyte (*Clonostachys rosea*). In terms of abundance, these ubiquitous OTUs accounted for 27% and 18% of the total reads for prokaryotes and fungi respectively. Particularly the peaks in distribution of OTUs over the samples differed between the prokaryotic and fungal community components. The prokaryotic distribution peaked around 8 samples (>200 OTUs), corresponding to the number of samples per site and suggesting an important role for locally-specific taxa in prokaryotic communities. On the hand, the fungal distribution peaked around 4 samples (>300 OTUs) and exponentially declined with occurrence, suggesting a role for both locally-specific and tree-specific taxa in the fungal communities.

**Figure 3.**
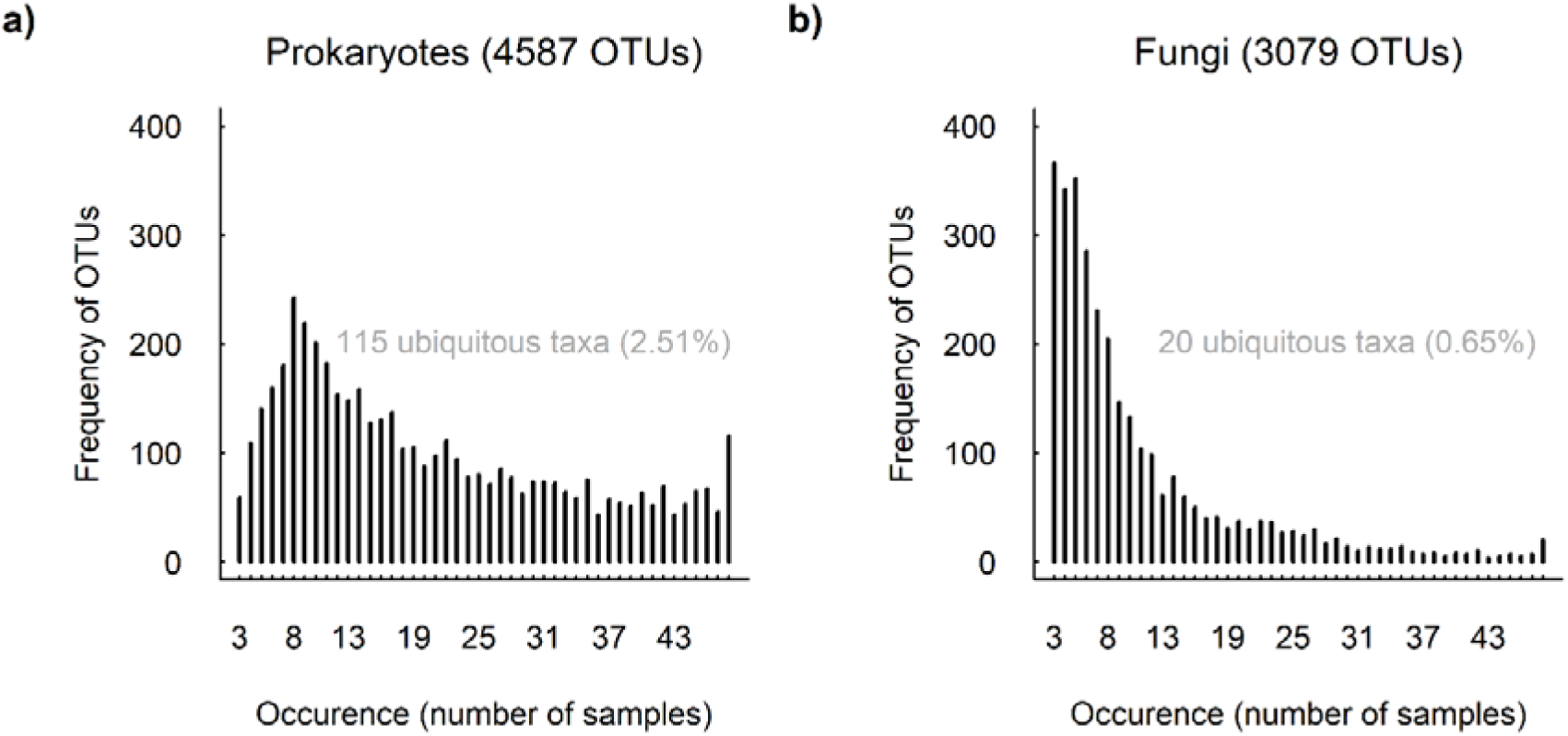
The prokaryotic **(a)** and fungal **(b)** frequency distribution of OTUs over the samples, where each OTU is assigned to the number of samples it occurs in. Taxa occurring in all samples (‘ubiquitous taxa’) are shown in gray, whereas most OTUs only occurred in fewer than 10 samples. Total number of OTUs per domain is indicated in brackets behind the title.

Both tree-specific prokaryotic and fungal taxa were found using logfold^2^ changes in abundance under *Q. ilex* versus *Q. suber* trees, independent of precipitation regime. Within prokaryotes, 61 taxa belonging to unique 44 families were significantly associated to one of the tree species (21 taxa to *Q. ilex*, 40 taxa to *Q. suber;* Figure 4a). Of those taxa, 23% belonged to *Chloroflexi*, 11% to *Alphaproteobacteria, Planctomycetes* and *Patescibacteria*, 10% to *Actinobacteria* and *Gammaproteobacteria*, 7% to *Verrucomicrobia*, and 2-3 % to each of the remaining phyla (*Firmicutes, Gemmatimonadetes, Cyanobacteria, Euryarchaeota, Acidobacteria*, WPS-2, *Bacteroidetes*). Even more fungi (126 taxa), belonging to 62 unique genera, were significantly associated to a particular tree species (72 taxa to *Q. ilex*, 54 taxa to *Q. suber*; Figure 4b). Of those taxa, 71% belonged to *Ascomycota*, 20% to *Basidiomycota*, 4% to *Chytridiomycota*, and 1% to each of the remaining phyla (*Mucoromycota, Rozellomycota Monoblepharomycota, Mortierellomycota*). Another 2% remained unclassified. The OTUs that could be classified to the genus level (75 of 126) mostly belonged to unique families and genera. Except within the *Ascomycota*, many families (*Herpotrichiellaceae, Coniochaetaceae Dothioraceae, Helotiaceae, Myxotrichaceae, Nectriaceae, Ophiostomataceae, Pyronemataceae, Sympoventuriaceae, Teratosphaeriaceae, Trichocomaceae*) harbored multiple tree-associated taxa, including mainly saprotrophic taxa. Within the Basidiomycota, four families (*Bulleraceae, Cortinariaceae, Russulaceae, Thelephoraceae*) harbored multiple tree-specific taxa, three of which were ectomycorrhizal. *Russulaceae* included the genera *Gymnomyces* and *Russula*. The *Thelephoraceae*, included *Pseudotomentella* and *Tomentella*, and the *Cortinariaceae* harbored more than one taxon (*Cortinarius* 4 OTUs). Finally, we found one family within the *Chytridiomycota* (*Spizellomycetaceae*), that harbored the tree-specific pathogenic genera (*Kochiomyces* (1 OTU) and *Spizellomyces* (2 OTUs).

**Figure 4.**
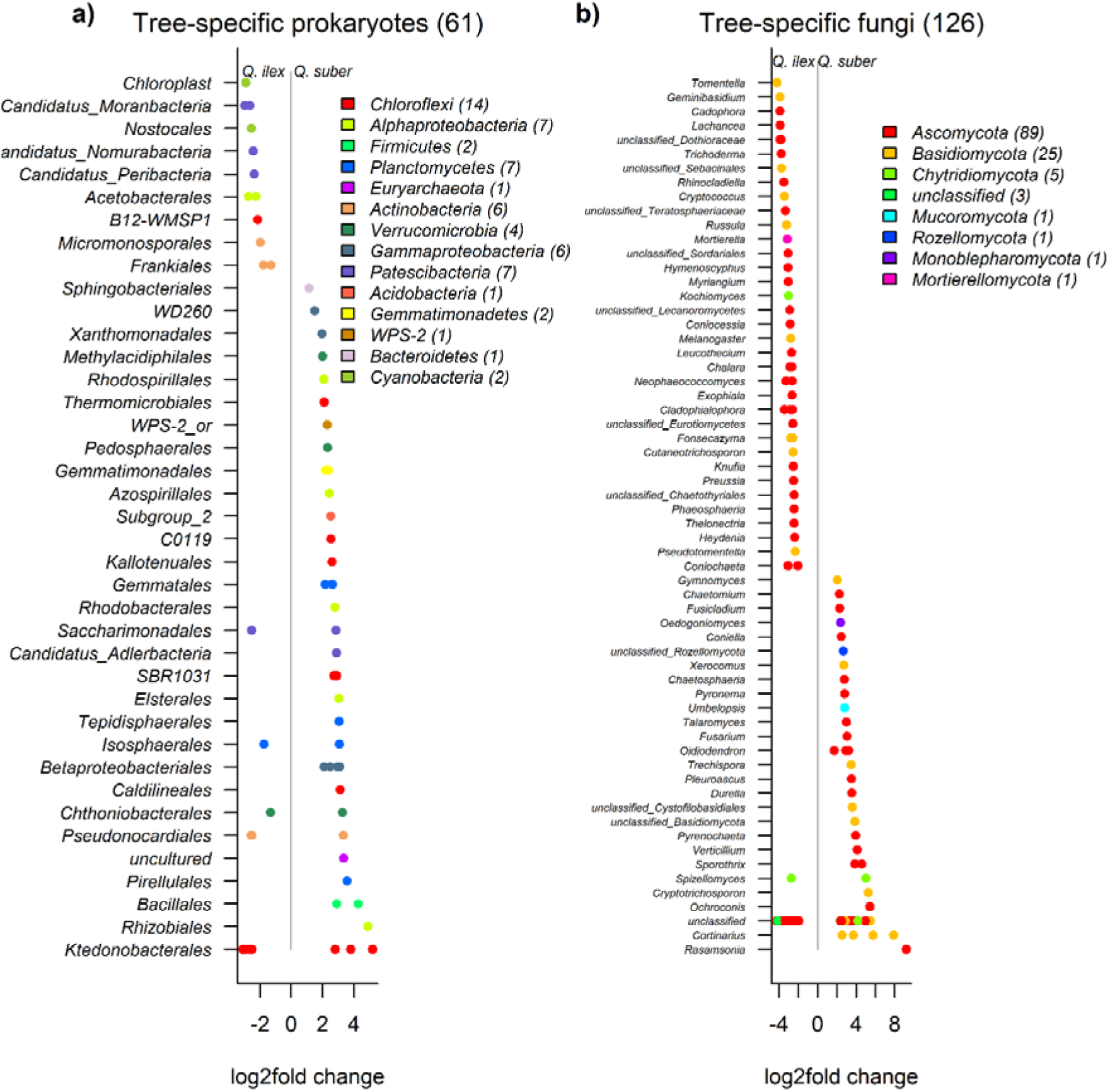
Tree-specific prokaryotic (a) and fungal (b) taxa, grouped by order (prokaryotes) and genus (fungi) and colored by phyla (legends). Log2fold change values indicate the strength and direction of the association to *Quercus ilex* (<0) and *Quercus suber* (>0). Significance (*P*<0.01) based on FDR corrected likelihood-ratio tests. Legends show phyla sorted by log2fold change abundance, with OTU richness per phyla given in brackets. Number of host-specific taxa per domain is indicated in brackets behind the title.

Locally-specific taxa, defined as taxa which were more abundant than expected based on random distributions over sites [38], were found in both prokaryotic and fungal communities (Figure S3). The proportion of locally-specific taxa for each site decreased along the precipitation gradient (prokaryotes and fungi analyzed together: F_1,8_=8.26, R^2^_adj_=0.32, *P*=0.021, Figure 5), indicating higher specialization rates for prokaryotes and fungi occurring at the drier end of the precipitation gradient.

**Figure 5.**
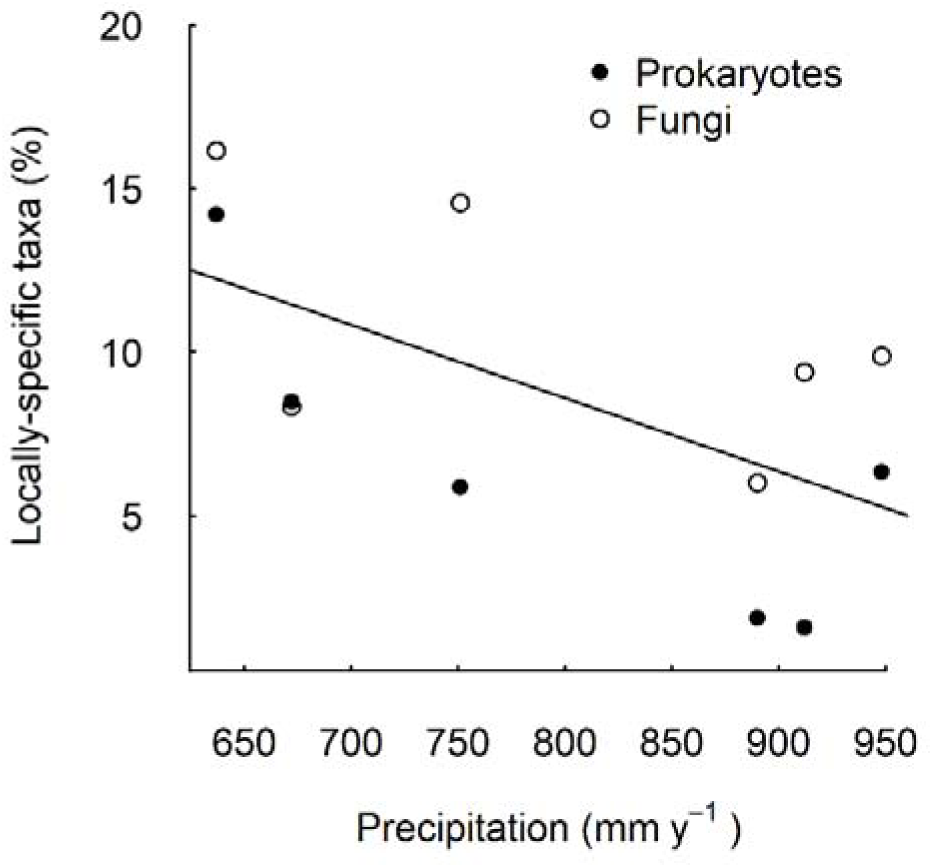
Proportion of locally-specific taxa for prokaryotes (filled circles) and fungi (open circles) along the precipitation gradient. More local indicator taxa were found in drier than in wetter areas. The trend line corresponds to the linear model including both community types (F_1,8_ = 8.26, R^2^_adj_ = 0.32, *P* = 0.021). Separate linear model for prokaryotes (F_1,4_ = 7.37, R^2^_adj_ = 0.56, *P* = 0.053) and for fungi (F_1,4_ = 2.27, R^2^_adj_ = 0.20, *P* = 0.20) respectively.

## Discussion

### Microbial community diversity under two oak species change along a precipitation gradient

Climate and precipitation in particular, can enhance overall prokaryotic and fungal diversity [12] and biomass [11]. In line with this, most microbial diversity measures showed positive trends along the precipitation gradient. However, fungal α diversity (richness) was not affected by precipitation, suggesting that precipitation has stronger effects on prokaryotes than fungi. Such differences in bacterial and fungal communities along precipitation gradients, suggest that fungi are more resistant to drier conditions than bacteria [39,40], however also opposing trends have been reported. Such differences could potentially be explained by the copiotroph–oligotroph trade-off, also known as r- and K-selection theory [17]. For example, Proteobacteria and Actinobacteria tended to be more ubiquitous in the nutrient-rich soils in the wetter soils. On the contrary, Acidobacteria are known to be ubiquitous, diverse, desiccation tolerant, and largely oligotrophic bacteria, abundant under low resource availability [41]. Yet, it is known that subgroups within a phylum might show different responses than the one observed at the phylum level, potentially leading to misinterpretations. For example, different subgroups in Proteobacteria and Actinobacteria might occupy different ecological niches and can show mixed responses to precipitation [20]. Former studies find fewer oligotrophs and more copiotrophs with increasing precipitation [17,20]. In our study, the relative abundances of the most abundant fungal phyla like *Ascomycota* and *Basidiomycota* did not vary significantly along the precipitation gradient. However, the relative abundance of the prokaryotic *Acidobacteria* increased along the precipitation gradient. Whereas changes in the relative abundances of *Patescibacteria* and *Mortierellomycota* along the precipitation gradient depended on tree species. Such opposing patterns between taxonomic groups have been reported before [10,42]. Notably, our results are in line with those from earlier studies focusing on mycorrhizal community changes in oaks across sites varying in water availability. For example, a long-term precipitation reduction experiment resulted in modified EMF communities under *Q. ilex* [43], and the mycorrhizal status of *Quercus agrifolia* shifted with soil moisture from predominantly EMF in wet to almost exclusively arbuscular mycorrhizal fungi (AMF) during severe droughts [44]. Here, we show that besides ectomycorrhizal fungi, also prokaryotic and fungal communities under several specific plant species vary along a precipitation gradient.

Host plant family and functional group determined the community composition of ectomycorrhizal fungi (EMF) in a global analysis [45] and in dry areas [46]. In Mediterranean ecosystems and temperate forests, even lower taxonomic levels (plant species identity) drove fungal richness and fungal community structure [47,48]. Despite the close phylogenetic relationship between the *Quercus* species, each species harbored distinct soil microbial communities. Moreover, we show that even within the functional group of evergreen oaks, differences in the relative abundance of *Basidiomycota* (which includes the EMF-rich *Agaricomycetes* class) can be found. This adds to studies comparing the EMF communities of evergreen and deciduous oaks [27,49]. It would be interesting to conduct further studies including a larger pool of congeneric tree species to better understand what root or leaf traits might drive this variation in soil communities. Particularly because it has been shown that such variation in the soil microbiome can affect the performance of seedlings growing under congeneric trees [36].

The setup of our study sites was intended to test for differences in the soil microbiome between oak species along a precipitation gradient, but it also included a natural variation in soil properties, possibly partly caused by the different tree species (e.g. litter input and root exudates). The majority of earlier studies along environmental gradients have acknowledged the importance of additional or indirect factors (e.g. edaphic conditions, soil properties and plant communities) in structuring the soil microbiome [10–12]. Our study confirmed this, and we found that soil calcium and pH were highly correlated and important variables for explaining variation in both prokaryotic and fungal communities. At the global scale, these same two variables emerged as important predictors of fungal richness [10], but pH did not drive patterns in prokaryotic communities in drylands [12]. Future studies and experiments are needed to better understand the separate effects of calcium and pH on soil geochemical processes.

In line with our expectations, edaphic conditions affected prokaryotes more than fungi. Notably, only prokaryotes were affected by variations in soil texture and ammonium, whereas only fungi were impacted by changes in soil organic matter content. Whereas fungi can spread their mycelium around, attaching to resource patches and plant roots, prokaryotes are more constrained to the soil matrix and pores, which makes it more likely for the latter to be affected by soil chemistry [17,28]. Taken together with differences in litter inputs and root exudates among trees [26,27], this finding might also explain why we found fungal diversity, in terms of richness and community structure, to be more related to tree species than prokaryotic communities. Related to this, with increasing precipitation we found greater phosphorus concentrations under *Q. suber*, whereas calcium concentrations decreased under *Q. ilex* along the precipitation gradient (Table S3). Long-term field experiments designed to separate precipitation from other tree-related processes, including litter input and root exudation, would be particularly valuable for exploring possible explanations in more detail.

### Specialized prokaryotes and fungi under two oak species along a precipitation gradient

Unravelling the individual contributions of soil microbial taxa facilitates an improved understanding of the role that interactions between plants and soil microbes play in shaping the responses of plant community composition and ecosystem processes to global environmental changes [50]. Soil microbial components including enemies, symbionts and decomposers, may show different responses resulting in different net magnitudes and directions of plant–soil feedbacks along various environmental gradients [3,6,51],. Plant-soil feedbacks require specific variations in the soil communities of a host species, leading to asymmetric fitness differences in subsequent plant growth [52]. Our study provides evidence for such mechanisms as we found proportion of ubiquitous taxa in the fungal communities (0.65%) as well as in the prokaryotic communities (2.46%), suggesting that not all microbes occur everywhere, but implying a certain degree of specialization, particularly within the fungal communities. Particularly, we found more fungal taxa (173 OTUs) than prokaryotic taxa (40 OTUs) associated with a specific tree species. None of the fungal genera harbored specialized taxa under both tree species, except for the saprotrophic genera *Cladophialophora* and *Cryptococcus* and the plant pathogens *Spizellomyces*. Even on a family level, we only identified two families that harbored plant-specific taxa for both oak species. First, some ectomycorrhizas within *Russulaceae* (genus *Lactarius* and *Gymnomyces*) were specifically associated with *Q. suber*, whereas others from the confamiliar genus *Russula* were associated with *Q. ilex*. Second, in the family *Spizellomycetaceae* (*Chytridiomycota*), the plant pathogens *Kochiomyces* and an unclassified genus were associated with *Q. ilex*, whereas the confamiliar *Spizellomyces* was associated with *Q. suber*. These results suggest that a fraction of the soil microbiome harbors associated fungi, including mycorrhiza, saprotrophs and pathotrophs that are specific to a plant species, even when the plants are closely related.

The effects of abiotic conditions on interactions between plants and their soil microbes remain unclear. For example, a reduced water availability has been shown to result in smaller net negative effects of the soil microbial community on conspecific plant performance [36,37]. This suggests either a decrease in specific antagonistic interactions, an increase in mutualistic interactions or a combination of both [51–53]. Most studies so far have generally found negative effects of the conspecific soil community on subsequent plant growth [2,52]. A test with savanna tree species found positive heterospecific feedbacks suggesting an important role for mutualists in drier environments [53]. Droughts can reducing plant-microbe associations and neutralize plant soil feedbacks [54] and inconsistent abiotic conditions between training phase and feedback phase resulted in variable feedback effects [55]. We found that microbial communities harbored more locally-specific taxa in the drier sites than in the wetter sites, and particularly locally-specific prokaryotes decreased with increasing precipitation. The occurrence of more specialized taxa at the dry end of our gradient, could be driven by facilitation, whereas more generalist taxa in more humid areas, are possibly driven by competition [3,36]. A similar pattern was observed for the locally-specific fungal taxa along the precipitation gradient, although no significant relationship was detected. It is possible that the difference between prokaryotic and fungal communities was found because fungal communities are more buffered by host plants whereas bacteria are more strongly dependent on edaphic conditions [17,28]. An additional explanation could be that fungi are more resistant to drier conditions than bacteria [22], reinforcing our first hypothesis. Our study serves as a proof of concept, which to our knowledge has not been demonstrated before. It is bringing us one step closer to identify potential crucial habitat-specific prokaryotes and fungi of particular ecological importance for plant-soil interactions along gradients.

## Conclusion

In the last years, several studies have discussed the climate-dependency of microbial communities, their interactions with and feedbacks on plants [55–57]. Here we show that plants can harbor distinct associated microbes when climatic conditions change. Particularly, drier conditions resulted in a higher proportion of locally-specific taxa as compared with soil communities at the wetter end of the precipitation gradient. Moreover, we show that besides frequently studied fungal associations such as mycorrhiza, also prokaryotes associate with tree species and might therefore potentially play a prominent role generating plant-soil feedbacks.

Our approach highlights taxa in the soil microbial community which are potentially important in generating precipitation depended differences between host plant species. Such approach goes beyond alpha and beta diversity measures of the microbiome and is not limited to taxonomic or functional knowledge of the taxa. This can help to discover microbial taxa which are important in generating context-dependent plant-soil feedbacks, which in turn help to better understand underlying mechanisms of ecosystem functioning and vegetation dynamics under global change.

## Acknowledgements

We thank P. Ruiz-Benito and L. Matías for help with third national forest inventory (NFI3) and field site selection. We are grateful to E. Gutiérrez-González and B. Stierli for their help in the laboratory, J. Domiguez, M. Renner, C. Saladin and A. Locher for help in the field, the Genetic Diversity Center (GDC) at ETH Zürich and the McGill University and Génome Québec Innovation Center in Montreal, Canada for performing Illumina MiSeq sequencing. M. Dawes for textual improvements and A. Frossard and T. Wubet for useful comments on earlier versions of the manuscript.

## Availability of data and material

The datasets generated and/or analyzed during the current study are available in the NCBI Sequence Read Archive repository, under the BioProject accession number PRJNA550539 [www.ncbi.nlm.nih.gov/bioproject/?term=PRJNA550539]

## Authors’ contributions

The study was initiated and performed by GR, who gained funding for the study. Laboratory analyses were facilitated by LGA and BF. BF conducted the bioinformatics. GR analyzed the resulting data and wrote the first draft, with the help of LGA and BF. All authors contributed to revisions of the manuscript.

