## supplementary for "Mediterranean oaks harbor more specific soil microbes at the dry end of a precipitation gradient"

### Mediterranean oaks harbor distinct soil microbiomes along a precipitation gradient

3Current address: German Centre for Integrative Biodiversity Research (iDiv) Halle-Jena-Leipzig, Deutscher Platz 5e, 04103, Leipzig, Germany. Martin-Luther-University Halle-Wittenberg, Institute of Biology/Geobotany and Botanical Garden, Am Kirchtor 1, 06108, Halle, Germany

**Supplement 1.** Extended methods for assessing soil physical and chemical properties

The Bouyoucos hydrometer method (Day, 1965) was used to assess soil texture. To assess soil pH, 10 g soil was dissolved in 25 ml Elix H_2_O and 10 g soil in 25 ml KCL (1 M). The tubes were shaken (30 min; 150 rpm, OVAN shaker) and centrifuged (4 min; 4000 rpm, KUBOTA 2010) before pH was measured (CRISON pH meter basic 20+). Organic carbon of 0.5­1 g of soil was assessed by the oxidation of organic matter with potassium dichromate and sulfuric acid and determined by spectrophotometry (600 nm), according to the Walkley-Black protocol (Walkey 1946). The soil was calcined in an oven muffle at 540°C to determine the total organic matter content (Anderson 1982). NO_3_^-^ and NH_4_^+^ were extracted from 1 g of soil with 1 M KCl and determined by spectrophotometry (550 nm). Another 1 g of soil was dissolved in 10 ml HCl NH_4_F according to the P-Bray method (Bray and Kurtz 1945) to assess available P. The solution was assessed at 880 nm in a spectrophotometer (filter-based multi-mode microplate reader: FLUOstar Omega; BMG LABTECH). Ca, Mg and K were extracted from 2 g of soil with 1 M ammonium acetate and determined by atomic emission spectroscopy (Pratt, 1965). All results were double-checked and repeated when needed. Correlations between variables were checked and highly correlated variables were omitted (correlation matrix not shown).

We found overall differences in soil properties between the tested tree species along the precipitation gradient, as indicated by a significant interaction between precipitation and tree species (Table S2). In particular, phosphorus increased under *Q. suber* along the precipitation gradient, whereas calcium decreased under *Q. ilex* along the precipitation gradient. Even though calcium and pH were correlated (F_1,46_=67.96; R^2^=0.57; P= <0.001), no significant changes in pH were found between the two species or along the precipitation gradient.

**Supplement 2. Characterization of the soil microbial communities**

A total of 4,934,489 16S and 3,013,047 ITS sequences were retrieved from sequencing, and a total of 1,043,273 (mean 21,735 ± SD 3674 per sample) prokaryotic and 1,981,289 (mean 41,277 ± SD 13,583 per sample) fungal high-quality sequences were kept after filtering. Sequence clustering yielded 4,587 prokaryotic and 3,079 fungal OTUs. A complete list of the detected unfiltered prokaryotic and fungal taxa, from phylum to OTU level, including abundance information, have been deposited as raw sequences in the NCBI Sequence Read Archive under the BioProject accession number PRJNA550539.

In brief, *Planctomycetes* (23.8%), *Alphaproteobacteria* (18.3%), *Verrucomicrobia* (11.7), *Acidobacteria* (11.3), *Actinobacteria* (11.1), *Chloroflexi* (7.1%), *Gammaproteobacteria* (4.6%), *Patescibacteria* (3.5%), Firmicutes (2.5%), *Bacteroidetes* (1.7%), *Deltaproteobacteria* (1.2%) were the most abundant prokaryotic phyla (Fig. S3). The ten most abundant prokaryotic genera were *WD2101* (6.4%), *Candidatus* *Udaeobacter* (7.1%), *Pir4_lineage* (3.3%), *Sphingomonas* (1.8%), *Bacillus* (1.8%), *Bryobacter* (1.8%), *Acidothermus* (1.8%)*.* Of the prokaryotic OTUs, 12.6% remained uncultured and 9% unclassified at the genus level.

For the fungi, *Ascomycota* (63.2%), *Basidiomycota* (31.2%), Mortiellomycota (2.2%) and the Mucoromycota (2.1%) were the most abundant phyla and *Chytridiomycota*, *Glomeromycora* and *Blastocladiomycota* each accounted for less than 1% of the total abundance (Fig. S3). *Penicillium* (7.7%), *Russula* (7.3%), *Cortinarius* (6.0%), *Cladophilalophora* (4.6%), *Inocybe* (3.3%), *Cenococcum* (2.5%), *Mortierella* (2.2%), *Humicola* (2.1%), *Trichoderma* (1.7%) and *Umbelopsis* (1.6%) were the ten most abundant fungal genera. Of the fungal OTUs, 17.7% remained unclassified at the genus level. To better understand the distributions of taxa among phyla, we assessed the relationship between taxa abundance and sample occurrence (prevalence) for each phylum (figures not shown).

**Table S1.** Summary results of post-hoc tests to assess which phyla responded to our treatments, using the ‘lmer’ function in the *lme4* package, with subsequent ANOVAs (all analyses completed in R). Significant terms shown in bold (P<0.1).

|  |  |  | Precipitation | | | Tree species | | |  | P X T | | |
| --- | --- | --- | --- | --- | --- | --- | --- | --- | --- | --- | --- | --- |
|  | **Prokaryotic phylum** | DF | Δ AIC | Chi Sq | P value | Δ AIC | Chi Sq | P value | DF | Δ AIC | Chi Sq | P value |
| 1 | Planctomycetes | 4,5 | -1.76 | 0.24 | 0.624 | -1.83 | 0.17 | 0.682 | 5,6 | -0.65 | 1.35 | 0.245 |
|  | α-proteobacteria | 4,5 | -1.99 | 0.01 | 0.929 | -1.55 | 0.45 | 0.502 | 5,6 | -1.72 | 0.28 | 0.595 |
|  | Verrucomicrobia | 4,5 | -1.98 | 0.02 | 0.881 | 1.67 | 3.67 | **0.055** | 5,6 | -1.17 | 0.83 | 0.363 |
|  | Acidobacteria | 4,5 | 0.83 | 2.83 | **0.092** | -2.00 | 0.00 | 0.964 | 5,6 | 2.61 | 4.61 | **0.032** |
|  | Actinobacteria | 4,5 | -1.26 | 0.74 | 0.389 | -1.06 | 0.94 | 0.332 | 5,6 | 0.18 | 2.18 | 0.140 |
|  | Chloroflexi | 4,5 | -1.14 | 0.86 | 0.354 | 0.29 | 2.29 | 0.130 | 5,6 | -2.00 | 0.00 | 1.000 |
|  | γ-proteobacteria | 4,5 | -2.00 | 0.00 | 0.953 | 1.47 | 3.47 | **0.062** | 5,6 | 0.52 | 2.52 | 0.113 |
|  | Patescibacteria | 4,5 | -1.08 | 0.92 | 0.338 | -1.77 | 0.23 | 0.635 | 5,6 | 2.35 | 4.35 | **0.037** |
|  | Firmicutes | 4,5 | -1.53 | 0.47 | 0.494 | -0.99 | 1.01 | 0.314 | 5,6 | -1.94 | 0.06 | 0.809 |
|  | Bacteroidetes | 4,5 | -1.74 | 0.26 | 0.613 | -1.28 | 0.72 | 0.397 | 5,6 | -1.37 | 0.63 | 0.428 |
|  | **Fungal phylum** | DF | Δ AIC | Chi Sq | P value | Δ AIC | Chi Sq | P value | DF | Δ AIC | Chi Sq | P value |
| 1 | Ascomycota | 4,5 | -1.20 | 0.80 | 0.372 | 0.98 | 2.98 | **0.084** | 5,6 | 0.71 | 2.71 | 0.100 |
|  | Basidiomycota | 4,5 | -1.36 | 0.64 | 0.425 | 1.84 | 3.84 | **0.050** | 5,6 | 0.38 | 2.38 | 0.123 |
|  | Mortierellomycota | 4,5 | -1.92 | 0.08 | 0.781 | -0.83 | 1.17 | 0.279 | 5,6 | 7.75 | 9.75 | **0.002** |
|  | Mucoromycota | 4,5 | -0.63 | 1.37 | 0.243 | -1.42 | 0.58 | 0.445 | 5,6 | -1.63 | 0.37 | 0.541 |
|  | unclassified | 4,5 | 0.63 | 2.63 | 0.105 | -0.94 | 1.06 | 0.303 | 5,6 | -0.08 | 1.92 | 0.166 |
|  | Chytridiomycota | 4,5 | -1.58 | 0.42 | 0.515 | -1.4 | 0.60 | 0.437 | 5,6 | -0.57 | 1.43 | 0.232 |
|  | Glomeromycota | 4,5 | -1.88 | 0.12 | 0.725 | -1.77 | 0.23 | 0.632 | 5,6 | -1.93 | 0.07 | 0.788 |

**Table S2.** Beta diversity of prokaryotic (left) and fungal (right) communities. Summary statistics based on PERMANOVAs with 9999 permutations, **a)** using Bray Curtis distances of the normalized abundances (Love et al 2014, McMurdie & Holmes 2014); **b)** using Euclidian distances of the square-root transformed relative abundances of the most abundant phyla. Significant terms (****P* < 0.001,** *P* < 0.01, **P* < 0.05,+ *P* <0.1) are shown in bold)

|  | Prokaryotic communities | | | | | |  | Fungal communities | | | | |
| --- | --- | --- | --- | --- | --- | --- | --- | --- | --- | --- | --- | --- |
| a) | DF | Sum Sq | Mean Sq | F_model_ | R^2^ |  |  | Sum Sq | Mean Sq | F_model_ | R^2^ |  |
| Precipitation (P) | 1 | 0.58 | 0.58 | 3.73 | **0.08** | **+** |  | 0.77 | 0.77 | 2.45 | **0.05** | ******* |
| Tree species (T) | 1 | 0.11 | 0.11 | 0.70 | 0.01 |  |  | 0.43 | 0.43 | 1.37 | **0.03** | ******* |
| S x T | 1 | 0.16 | 0.16 | 1.05 | **0.02** | **+** |  | 0.41 | 0.41 | 1.29 | **0.03** | ******* |
| Residuals | 44 | 6.86 | 0.16 |  | 0.89 |  |  | 13.89 | 0.32 |  | 0.90 |  |
| Total | 47 | 7.72 |  |  |  |  |  | 15.50 |  |  |  |  |
| b) | DF | Sum Sq | Mean Sq | F_model_ | R^2^ |  |  | Sum Sq | Mean Sq | F_model_ | R^2^ |  |
| Precipitation (P) | 1 | 0.06 | 0.06 | 1.57 | 0.03 |  |  | 0.06 | 0.06 | 1.57 | **0.03** | ***** |
| Tree species (T) | 1 | 0.12 | 0.12 | 2.82 | **0.06** | + |  | 0.12 | 0.12 | 2.82 | **0.06** | **+** |
| S x T | 1 | 0.08 | 0.08 | 1.98 | 0.04 |  |  | 0.08 | 0.08 | 1.98 | 0.04 |  |
| Residuals | 44 | 1.80 | 0.04 |  | 0.87 |  |  | 1.80 | 0.04 |  | 0.87 |  |
| Total | 47 | 2.06 |  |  | 1.00 |  |  | 2.06 |  |  | 1.00 |  |

**Table S3.** Differences in soil properties across the precipitation gradient and between the two tree species. Summary statistics based on PERMANOVAs on Euclidean distances of the scaled soil properties, with precipitation (P) and tree species (T) as fixed factors. The model was run with 999 permutations, and site was included in the strata to account for site effects. Lower part of the table shows post hoc tests for phosphorus and calcium concentrations separately. Significant terms are shown in bold (P<0.1).

|  | | DF | Sum Sq | Mean Sq | | F model | R^2^ | | P value | |
| --- | --- | --- | --- | --- | --- | --- | --- | --- | --- | --- |
| Precipitation (P) | | 1 | 48.51 | 48.51 | | 5.29 | 0.10 | | **0.057** | |
| Tree species (T) | | 1 | 5.30 | 5.30 | | 0.58 | 0.01 | | 0.313 | |
| P x T | | 1 | 12.75 | 12.75 | | 1.39 | 0.03 | | **0.031** | |
| Residuals | | 44 | 403.45 | 9.17 | | 0.86 |  | |  | |
| Total | | 47 | 470.00 | 1.00 | |  |  | |  | |
| **Soil variable** | | | | | DF | ΔAIC | | Chi Sq | | P value |
| Phosphorous | Precipitation (P) | | | | 4,5 | 1.75 | | 3.75 | | **0.053** |
|  | Tree species (T) | | | | 4,5 | 0.80 | | 2.80 | | **0.095** |
|  | P x T | | | | 5,6 | 5.19 | | 7.19 | | **0.007** |
| Calcium | Precipitation (P) | | | | 4,5 | -1.27 | | 0.73 | | 0.393 |
|  | Tree species (T) | | | | 4,5 | 0.76 | | 2.76 | | **0.096** |
|  | P x T | | | | 5,6 | 4.65 | | 6.65 | | **0.010** |

**Figure S1.** The relative abundances of OTUs over the most abundant phyla for **(a)** prokaryotic and **(b)** fungal communities. Bars represent the average abundances for each site × tree species combination (n=4).
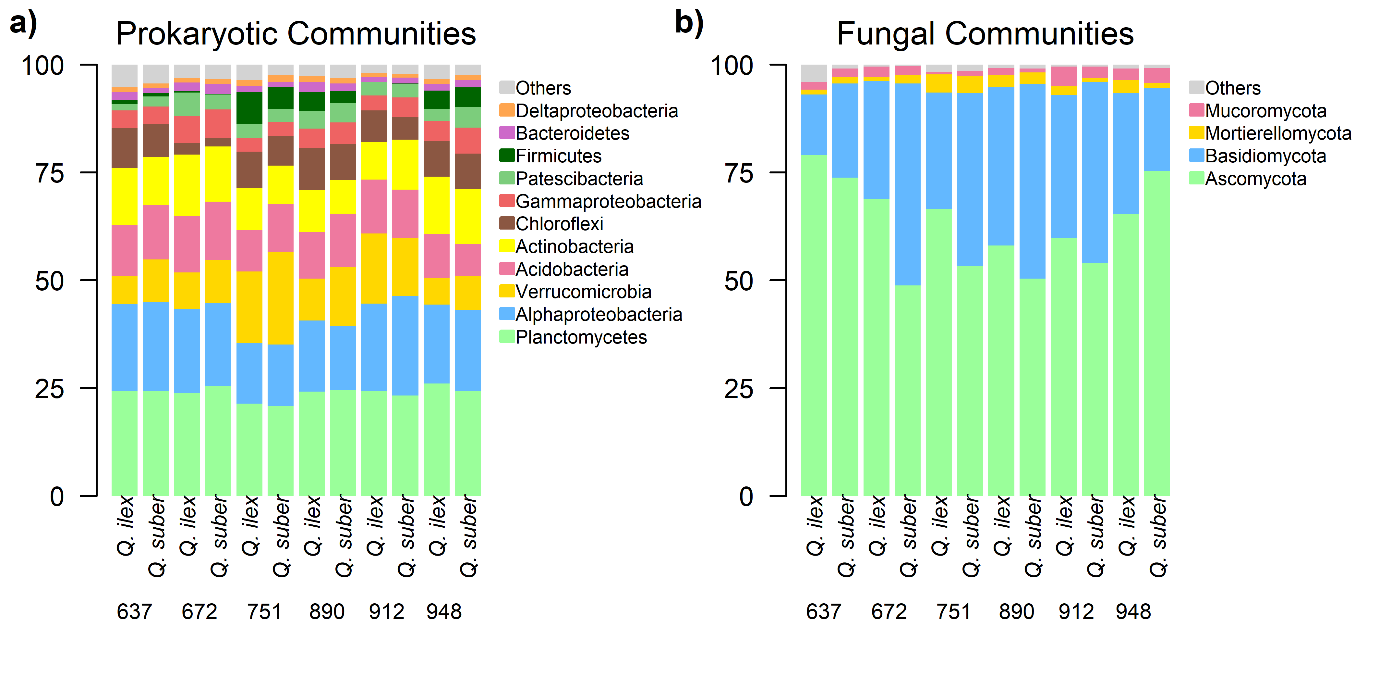

**Figure S2.** Rare faction curves of the prokaryotic (left) and fungal (right) communities. Curves represent samples colored by sampling site. Grey lines indicate minimum sample sizes to which all samples were rarefied prior to testing the effect of sample type on observed species richness and inversed Simpson index for **(a)** prokaryotic (15,548 reads per sample) and **(b)** fungal communities (12,740 reads per sample).

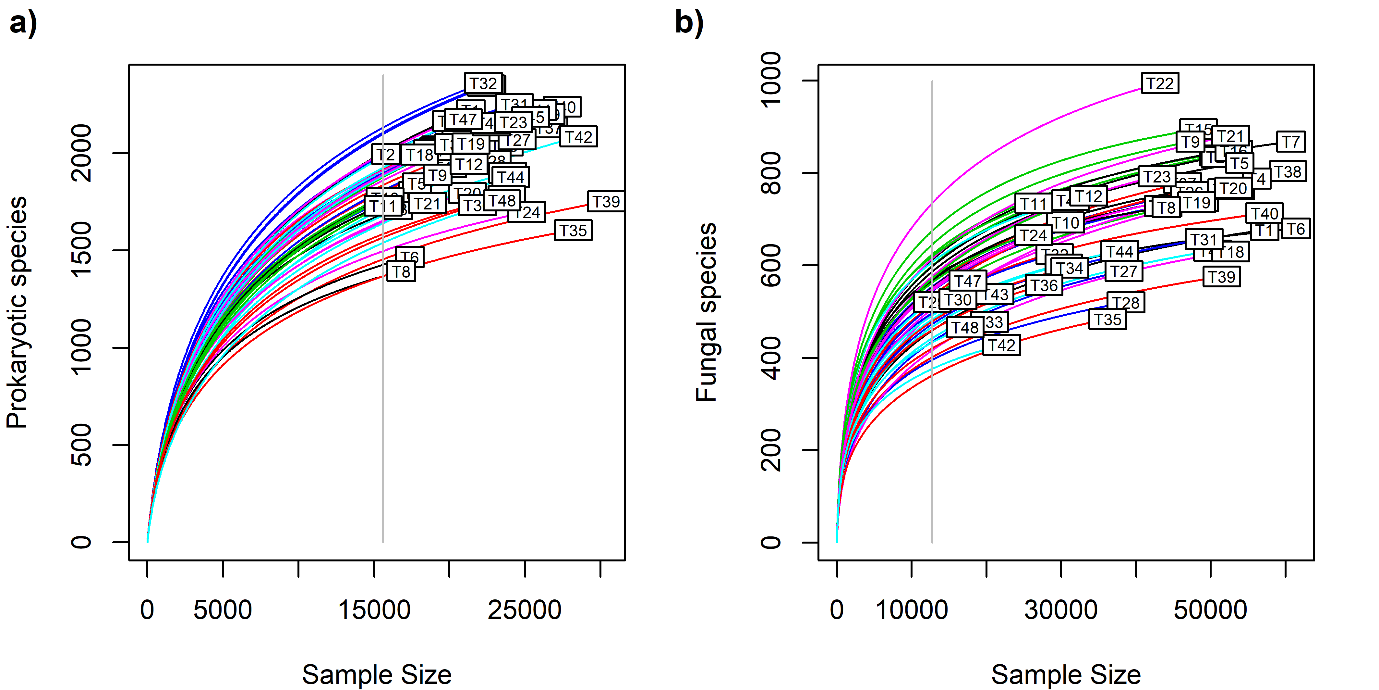

**Figure S3.** Local specificity for each of the sites ranging in precipitation (637, 672, 751, 890, 912 and 948 mm year^–1^). Expected local specificity under the null distribution where samples are randomly assigned to locations along the precipitation gradient (gray). Curves are loess regressions of the original (green) and randomized (gray) specificity values against abundance in the location, with associated confidence intervals. The upper six panels show **a)** prokaryotic communities and the lower six panels show **b)** fungal communities.

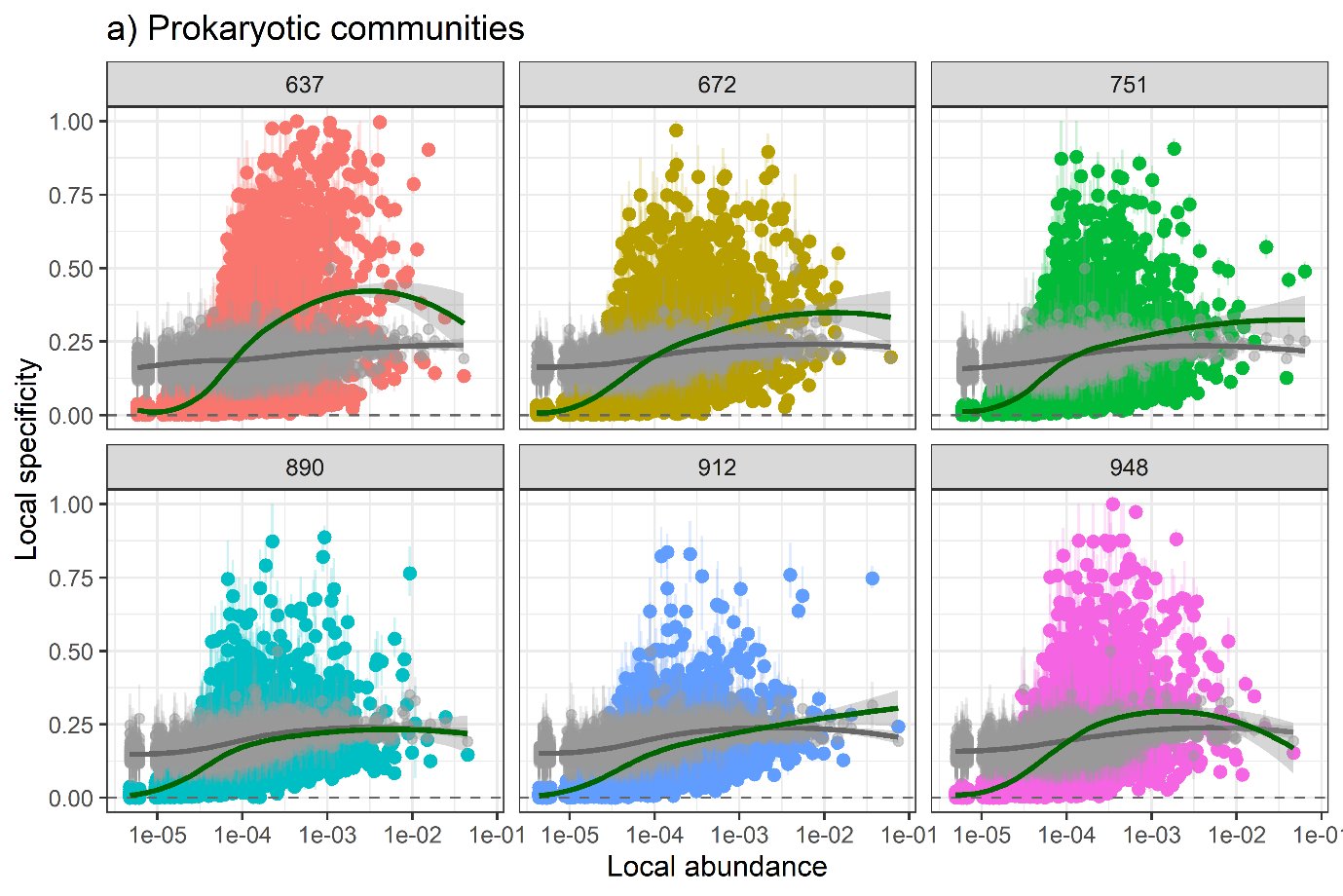

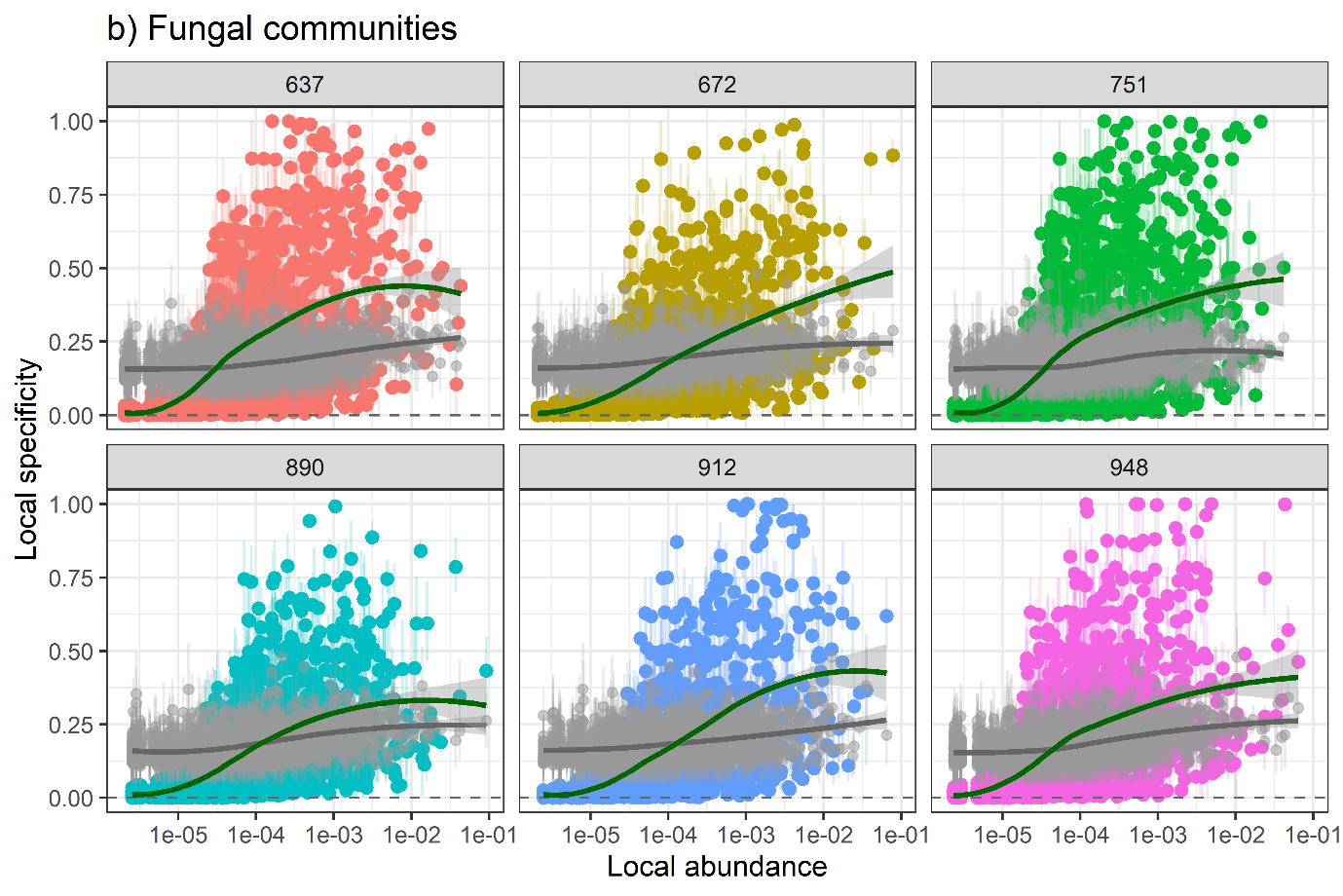
